# The Open Field test as a tool for behavior analysis in pigs - is a standardization of setup necessary? A systematic review

**DOI:** 10.1101/2021.09.27.461927

**Authors:** M. Schulz, L. Zieglowski, M. Kopaczka, R.H. Tolba

## Abstract

The Open Field test is a common tool to measure anxiety and behavioral changes in rodents. However, scientific findings of rodent experiments may not translate adequately to humans and it has been shown that larger animal models might perform better in that regard. As a result, the number of published studies involving the Open Field test in domestic pig models is increasing. The aim of our review was to investigate the Open Field set-ups in published studies as well as similarity between performance and parameters published. Following the PRISMA guidelines for reviews we selected 69 studies for data extraction in this systematic review. We were able to determine specific set-up conditions such as size, duration and daytime for most of the included studies and found a high variability within these test specifiers. Results indicate a non-uniform performance of set-up including size, timing, parameters and additional combined tests such as the novel object test. We would like to point out the need for standardization of Open Field test for pigs in order to improve result, comparability and reduce inconsistencies.

## Introduction

One tool to assess treatment outcomes is the Open Field (OF) test, which is widely used in rodent models. A closer look into history reveals that the OF was first developed by Hall in 1934 defined as an ‘unfamiliar enclosure’ [1]. It was supposed to analyze anxiety, behavior or toxicological effects in rodents. In the context of approach-avoidance conflicts the OF test however is recommended to be used in combination with other anxiety-related tests like elevated plus maze or the social interaction test in order to make a powerful statement, as OF alone is no valid indicator of emotions [2]. Within time and with evolving technology the OF was used to also determine distance and velocity as objective parameters for locomotor activity [3, 4]. Later, behavioral patterns such as rearing and grooming were added to determine well-being in rodents as they indicate stress levels [5–7]. In pigs, the OF is often combined with a <u>n</u>ovel <u>o</u>bject <u>t</u>est (NOT) or human approach test (HAT) [8]. For NOT an unfamiliar (new) object is usually hung from the ceiling, mostly after the pig adapted to the Open Field for a few minutes to assess the exploratory reaction. For the HAT, a human unfamiliar to the pig, enters the Open Field area and likewise the behavior such as anxiety or curiosity as well as the approach are recorded.

The domestic pig is a commonly used model in several fields of biomedical research [9]. It also closes the gap between rodents to humans with more genetically and morphological similarities to humans [10]. Especially before adapting clinical studies, a large animal model can answer specific questions in fields of surgical models, toxicology or brain disorder. However, behavioral tests established in rodents do not necessarily translate to pigs due to their different response to behavioral stimuli [11].

However, there are only a few validated methods to assess aversion to novelty in pigs [12]. In addition, animal models of pain-related studies include a variety of methods to assess changes in behavior following pain. Although, using rodents and flight animals for pain-assessment is criticized due to a lack of translational ability to humans [13–15]. Additionally, toxicology studies in pigs have also been used to investigate fetal programming, [16] antipsychotic drugs[17, 18] for mental disorders like schizophrenia or addiction to drugs [19]. Furthermore, environmental enrichment is known to have a positive influence on early life stage in regard of sensory, social, cognitive and motor functions in rats [20, 21]. It is also under investigation if such factors facilitate social or emotional behavior in pigs by rearing the environment. Especially in respect to farm animals the OF might be a powerful tool to determine the impact of husbandry conditions on pigs [22–24].

According to Forkman et al when transferring the OF test to domestic and farm animals their species specific behavior was not taking into account [25]. Rodents naturally show thigmotaxis when exposed in the OF whereas such behavior is not described for farm animals like pigs. Although boars could be seen roaming close to timberline, the distance to trees is with an average of 54m too high to be seen as thigmotaxic behavior and cannot be transferred to the OF arena [8]. Therefore, using OF as an pure indicator [24] of anxiety should be seen critical [25].

A screening of literature yields a wide and non-uniform use of the OF test in pigs varying in size, duration and outcome parameters. Additionally, a lack of information regarding the housing conditions, animals and performance of behavioral tests could be observed. Additionally, since there is so far no consensus on the behavioral parameters with the most significance for the emotionally state, impacts on social isolation or pain-based behavior change, we examined the broad range of parameters used in the studies to see which ones where the most often used.

Therefore, this systematic review investigates if an overlap of set-up, performance and parameters can be found between the studies to give future recommendations on standardization of OF test in pigs.

## Methods

This study was done in accordance with the guidelines of Preferred Reporting Items for Systematic Reviews and Meta-Analyses (PRISMA) [26] and was conducted using a registered protocol of the International Prospective Register for Systematic Reviews (PROSPERO, CRD42019156734).

### Search strategy

The survey aimed to identify published manuscripts with defined set-ups using OF in combination with *sus scrofa*. We searched MEDLINE (Pubmed), EMBASE and Web of Science with various search terms related to pig namely: “pig”, “porcine”, “swine”, “boar”, “gilt”, “miniature swine”, “sows”, “piglet” and “piglets” and combined them with terms to investigate into studies using the OF test to assess behavior with the terms “open field”, “open-field” and “behav*”. Search terms were similar for each database. The systematic literature search of studies was carried out in November 2019 for the years 1967 to November 2019.

### Study selection

We defined the inclusion criteria *a priori* into 4 dimensions: (I) a full text could be retrieved, (II) the study was written in English, (III) the species was limited to *sus scrofa* and (IV) the study used an OF with or without a novel object (NOT) or human approach test (HAT). We defined the OF as an unfamiliar, confined environment including alternative names like “pen” or “home pen” if used.

Studies were excluded if (I) the publication type was a review and (II) if studies were unpublished yet. All references were imported to Endnote X9 (Clarivate, Pennsylvania) and duplicates were removed.

### Primary Outcomes

- Set-up of OF (dimension, segments, duration, daytime)
- Age or body weight of animals

### Secondary Outcomes

- Time spent with NOT, HAT or intruder
- Parameters gained by OF (e.g. mobility, vocalization, exploratory behavior)
- Categorization of scientific fields of selected studies

### Data Collection

Two independent reviewers evaluated with a standardized selection worksheet the eligibility of each article. Studies were excluded when both reviewers agreed that the inclusion criteria were not met. A third reviewer was not required as all disagreements could be solved by discussion.

### Risk of Bias

The risk of bias was described and judged according to the SYRCLE’s risk of bias tool [27] adapted from the Cochrane Risk of Bias (RoB) tool [28]. Studies will be categorized as low, high or unclear risk of bias in all ten domains. Three independent authors judged the RoB for each study.

### Data synthesis and statistical analysis

Our primary outcome was study parameters regarding the dimension of OFs as well as the actual performance (time used, acclimatization, NOT and HAT) of the behavioral tests. Additionally, parameters of interest occurred as secondary outcomes describing the day time of test performance, age or body weight of animals used, and the investigated behaviors of pigs. Additionally, the domains of scientific questions were evaluated. A meta-analysis was not performed due to a solely narrative description of data. To correlate parameters the software Graph Pad Prism 7.0 (Dan Diego, USA) was used with Pearson r and 95% confidence interval. Boxplot were given with whiskers displaying the 5-95% confidence interval and outliers. Results are given in mean or median with standard deviation. Scientific domains were displayed in a radar plot by visual-paradigm.com.

## Results

The initial search of each data base yielded a cumulative 379 hits. Both independent reviewers had a high level of agreement in which 11 studies had to be discussed. The abstract screening provided 81 studies. A second consensus meeting with full congruence excluded another 12 studies due to exclusion criteria. In total, a sum of 69 studies were included into this survey. Figure 1 displays the review process stepwise.

**Figure 1.**
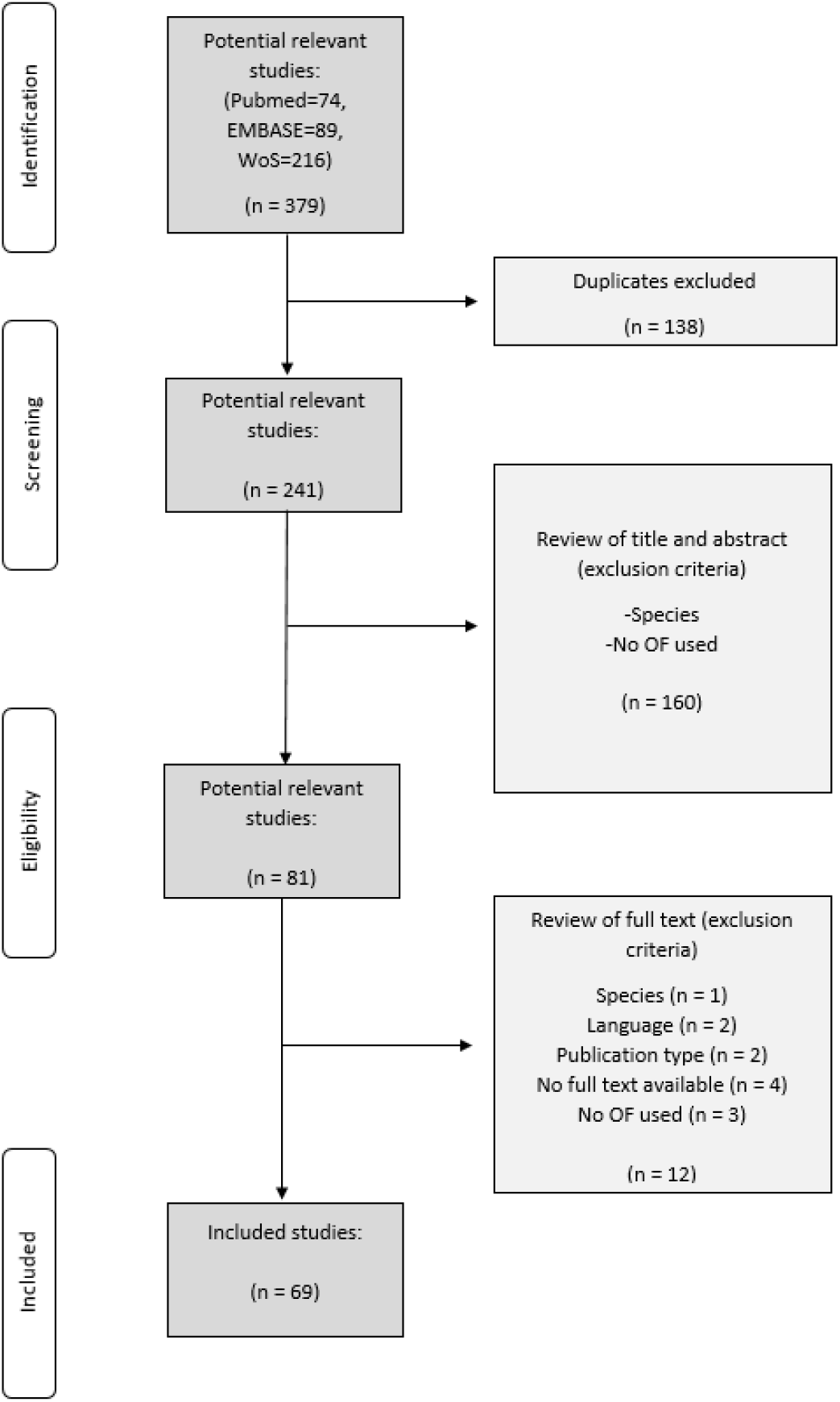
Flow chart of systematic review process by four stages according to the PRISMA guidelines

Our quality indicator between the 69 selected studies varied widely. Key criteria focused on detailed information given about OF dimension, day time of experiment, animals, test specifiers (time in OF, number of tests, etc.) and ethical approval by authorities.

Table 1 Quality and ranking of studies based on the given information sorts all studies by information available ranking those with highest quality above.

**Table 1.** Quality and ranking of studies based on the given information

| Author | Year | OF | Day time | Animals | Approved | Test specifiers | Sum per study |
| --- | --- | --- | --- | --- | --- | --- | --- |
| Donald [29] | 2011 | 1 | 1 | 1 | 1 | 1 | 5 |
| Lind [31] | 2005a | 1 | 1 | 1 | 1 | 1 |  |
| Von Borell [33] | 1992 | 1 | 1 | 1 | 1 | 1 |  |
| Bernardino [35] | 2016 | 1 | 1 | 1 |  | 1 | 4 |
| Castel [37] | 2018 | 1 |  | 1 | 1 | 1 |  |
| Clouard [39] | 2016 | 1 |  | 1 | 1 | 1 |  |
| de Oliveira [41] | 2015 | 1 |  | 1 | 1 | 1 |  |
| Fàbrega [43] | 2004 | 1 | 1 | 1 |  | 1 |  |
| Fijn [23] | 2016 | 1 |  | 1 | 1 | 1 |  |
| Gielsing [46] | 2014 | 1 |  | 1 | 1 | 1 |  |
| Herskin [48] | 2002 | 1 | 1 | 1 |  | 1 |  |
| Holme Nielsen [50] | 2018 | 1 |  | 1 | 1 | 1 |  |
| Horback [52] | 2018 | 1 |  | 1 | 1 | 1 |  |
| Jones [24] | 1998 | 1 | 1 | 1 |  | 1 |  |
| Kanitz [55] | 2004 |  | 1 | 1 | 1 | 1 |  |
| Laferriere [57] | 1995 | 1 | 1 | 1 |  | 1 |  |
| Leliveld [59] | 2017 | 1 |  | 1 | 1 | 1 |  |
| Meijer [61] | 2015 | 1 |  | 1 | 1 | 1 |  |
| Mosnier [63] | 2009 | 1 |  | 1 | 1 | 1 |  |
| Otten [65] | 2007 |  | 1 | 1 | 1 | 1 |  |
| Pond [67] | 2008 | 1 | 1 |  | 1 | 1 |  |
| Ralph [68] | 2018 | 1 |  | 1 | 1 | 1 |  |
| Siegford [70] | 2008 | 1 | 1 |  | 1 | 1 |  |
| Sullivan [72] | 2013-10 | 1 |  | 1 | 1 | 1 |  |
| Sumner [74] | 2008 |  | 1 | 1 | 1 | 1 |  |
| Thodberg [76] | 1999 | 1 | 1 | 1 |  | 1 |  |
| Zhong [78] | 2017 | 1 |  | 1 | 1 | 1 |  |
| Andersen [12] | 2000 | 1 |  | 1 |  | 1 | 3 |
| Andersen [81] | 2000 | 1 |  | 1 |  | 1 |  |
| Beattie [22] | 1995 | 1 | 1 | 1 |  |  |  |
| Brajon [84] | 2016 |  |  | 1 | 1 | 1 |  |
| Clouard [86] | 2015 | 1 |  |  | 1 | 1 |  |
| Dalmau [88] | 2009 |  | 1 |  | 1 | 1 |  |
| Dexter [90] | 1980 | 1 | 1 |  |  | 1 |  |
| Giroux [30] | 2000 |  |  | 1 | 1 | 1 |  |
| Giroux [30] | 2000 |  |  | 1 | 1 | 1 | 3 |
| Hessing [32] | 1994 | 1 |  | 1 |  | 1 |  |
| Horback [34] | 2016 | 1 |  |  | 1 | 1 |  |
| Jensen [36] | 1995 | 1 |  | 1 |  | 1 |  |
| Jones [38] | 2000 | 1 |  | 1 |  | 1 |  |
| Kanitz [40] | 2014 |  |  | 1 | 1 | 1 |  |
| Kanitz [42] | 2009 |  |  | 1 | 1 | 1 |  |
| Kanitz [44] | 2019 |  |  | 1 | 1 | 1 |  |
| Kinder [45] | 2019 |  |  | 1 | 1 | 1 |  |
| Kulikov [47] | 2014 | 1 | 1 |  |  | 1 |  |
| Lind [49] | 2005b | 1 |  | 1 | 1 |  |  |
| Loijens [51] | 2002 | 1 | 1 | 1 |  | 1 |  |
| Magnani [53] | 2012 | 1 |  | 1 |  | 1 |  |
| Meunier-Salaün [54] | 1991 | 1 |  | 1 |  | 1 |  |
| Pearce [56] | 1993 | 1 |  | 1 |  | 1 |  |
| Rutherford [58] | 2012 |  |  | 1 | 1 | 1 |  |
| Rutherford [60] | 2006 |  |  | 1 | 1 | 1 |  |
| Spoolder [62] | 1996 | 1 |  | 1 |  | 1 |  |
| Stracke [64] | 2017-02 |  |  | 1 | 1 | 1 |  |
| Stracke [66] | 2017-04 |  |  | 1 | 1 | 1 |  |
| Sullivan [11] | 2013-04 | 1 |  |  | 1 | 1 |  |
| Taylor [69] | 1986 |  | 1 | 1 |  | 1 |  |
| Val-Laillet [71] | 2013 |  | 1 | 1 | 1 |  |  |
| van der Staay [73] | 2009 |  |  | 1 | 1 | 1 |  |
| Von Borell [75] | 1991 | 1 | 1 |  |  | 1 |  |
| Weaver [77] | 2000 | 1 | 1 | 1 |  |  |  |
| Zebunke [79] | 2013 |  |  | 1 | 1 | 1 |  |
| Amory [80] | 2000 | 1 |  | 1 |  |  | 2 |
| Beilharz [82] | 1967 |  |  | 1 |  | 1 |  |
| Puppe [83] | 2007 |  |  | 1 |  | 1 |  |
| Taylor [85] | 1987 |  | 1 | 1 |  |  |  |
| von Borell [87] | 2009 |  |  | 1 |  | 1 |  |
| Zebunke [89] | 2017 |  |  | 1 |  | 1 |  |
| Puppe [91] | 1999 |  |  | 1 |  |  | 1 |
| Scott [92] | 2009 |  |  |  |  | 1 |  |

Only three studies were able to provide all information followed by 25 studies matching at least 4 of 5 quality criteria. More than half of the selected publications (41 publications) did provide 3 or less of the quality criteria and therefore had a lack of information.

### Set-up of Open Field

The size of the OF varied highly within the studies with a mean of 11.28 m^2^ and a large standard deviation of 8.55 m^2^. Two studies did not provide information about the size of their OF [30, 71] (Figure 2). Whereas most studies used a rectangular room, two studies of the same group mentioned a circular arena as OF [12, 81]. Forty-two studies gave additional information about a segmentation of the arenas in sections whether physically or virtual applied (Figure 3). The median of sections used is 12 ± 13.2.

**Figure 2:**
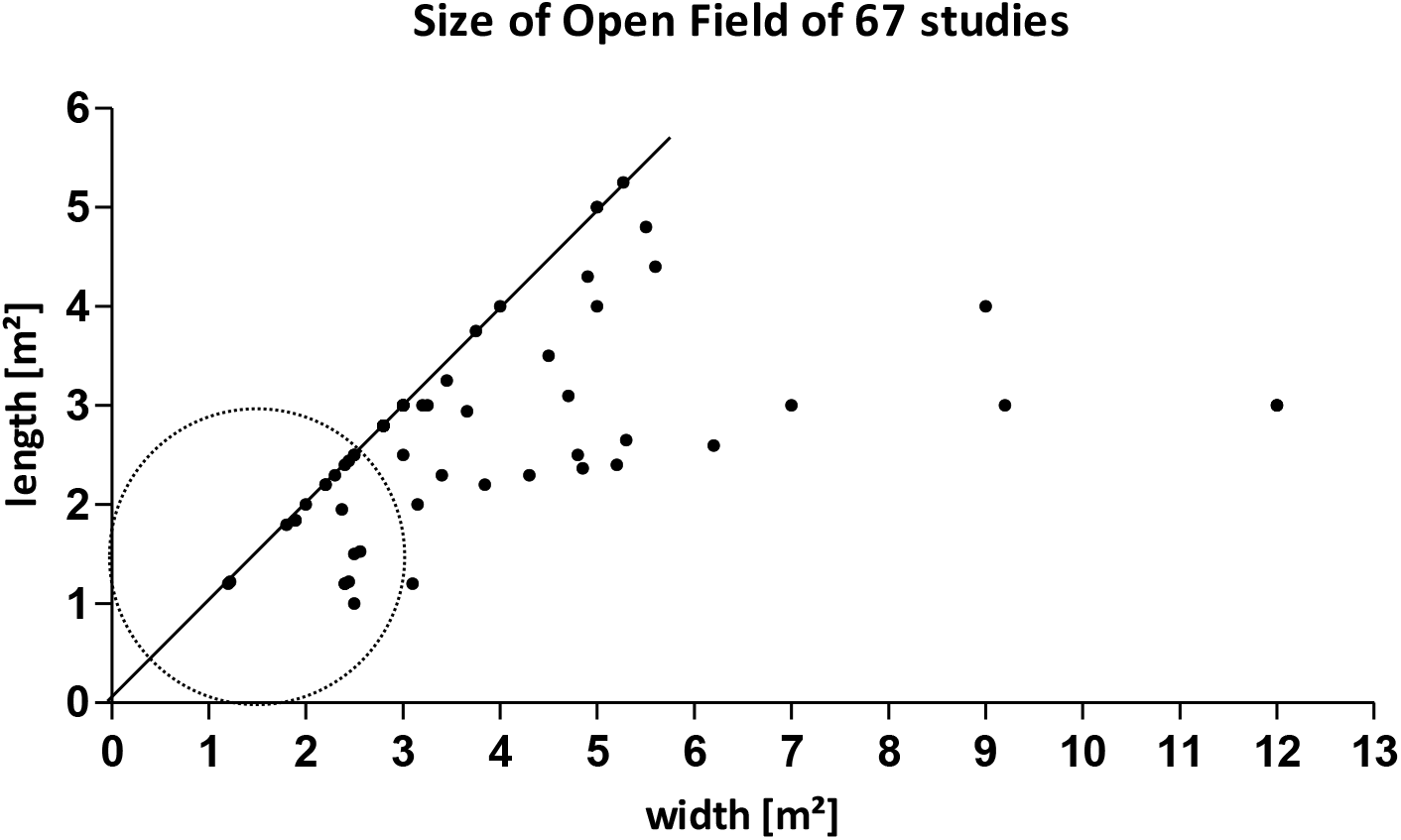
Points showing the length and width of OF with 27 points presenting a square (n=67). Two studies used a circular area which is presented as a circular dotted line.

**Figure 3.**
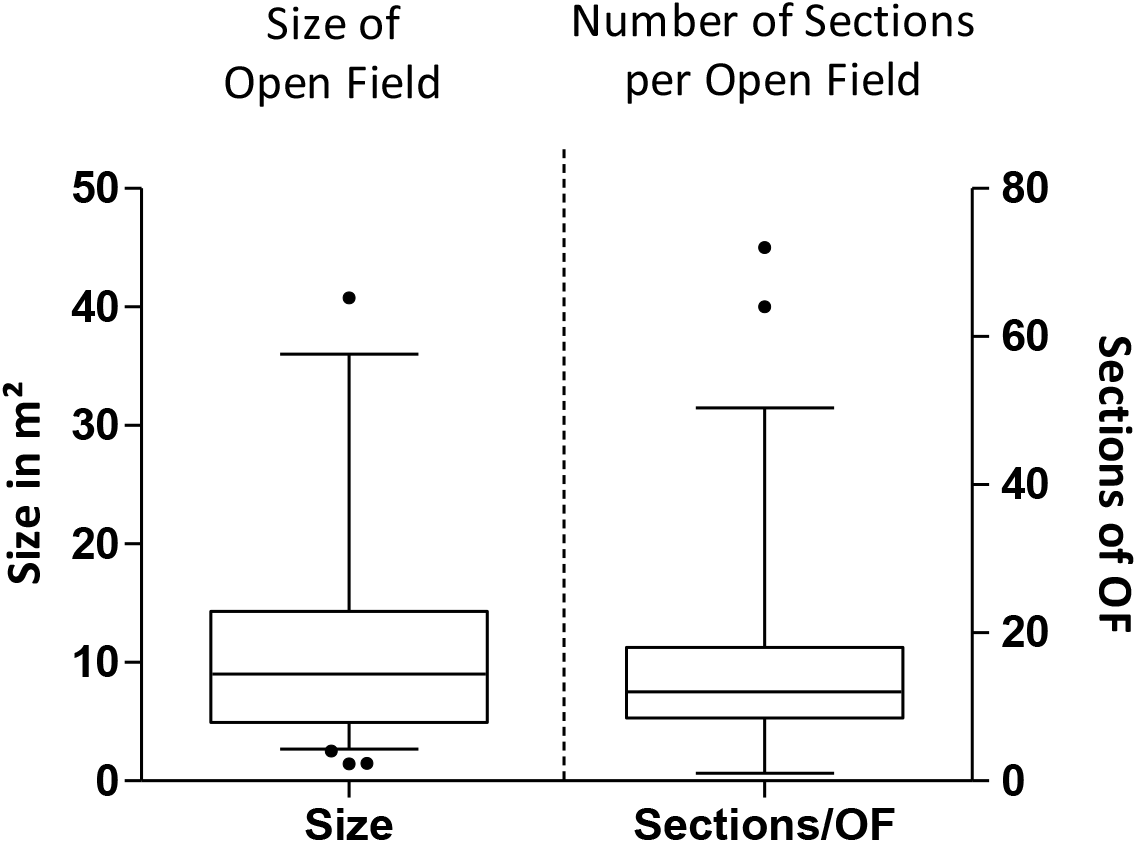
Left: Boxplot of Open Field size with whiskers displaying the 5-95% confidence interval and outliers (n=69). Right: Boxplot of number of segments used in the Open Field with whiskers displaying 5-95% confidence interval and outliers (n=45).

Some studies gave information about the size of the OF being adapted to the age of the pigs according to previous studies [12, 81]. To investigate, whether the size of OF correlates with the size of the animals, we looked into the weight and age distribution and compared them with the OF square meters. Overall, 6 studies did not provide information about age/body weight or size of OF and could not be included into the comparison [34, 37, 63, 71, 75, 92]. Even though most studies reported the age of their animals, the race and crossbreds varied, so no estimation can be made between age and body weight or size of the animal. As data of body weight are not uniform (given absolute [33, 54, 60, 62, 72, 93], as range [31, 47–50, 78], with mean and SD [12, 41, 51, 63, 80, 86, 88] or gained bodyweight due to food intake over time [44, 59]) a calculation was not possible. Thus, leading to 60 studies where information about the animals age could be included. For comparison, all ages were recalculated into days.

The distribution of age in the studies showed 63.8% (44 studies) of the animals being in the age of 2 to 79 days of which 13 studies used piglets during the weaning phase of <20 days [11, 30, 40, 42, 50, 55, 57, 65, 67, 71, 72, 90, 91]. Thirty-six percent reached an age of 80 days up to 540 days (Figure 4: Age distribution of animals per study. Age is given in days with a split x-axis at 100 days.). In. Six studies used adult pigs with an age over 6 months. [31, 49, 51, 77, 78, 85]

**Figure 4:**
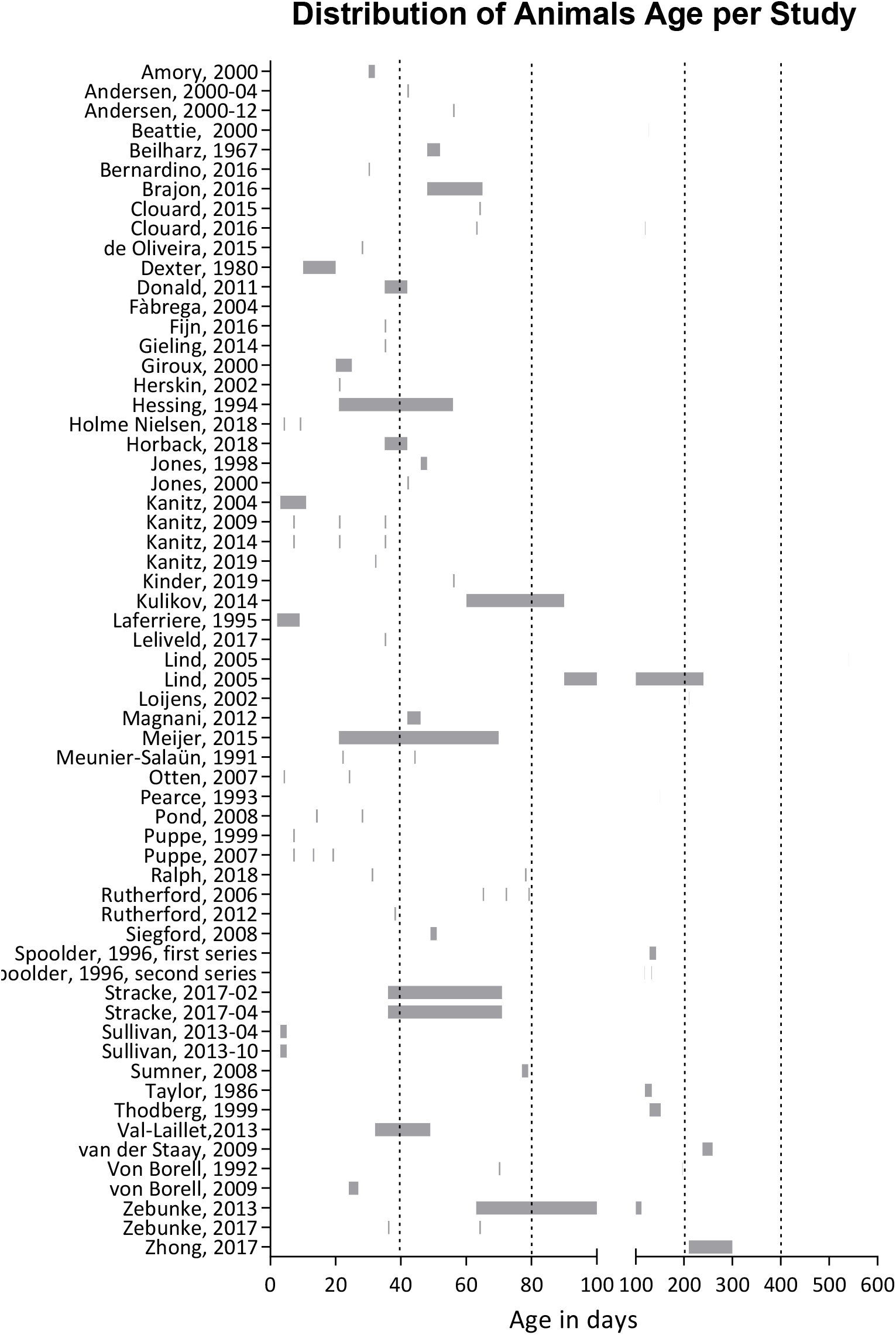
Age distribution of animals per study. Age is given in days with a split x-axis at 100 days.

The correlation of animals’ age against the size of OF showed no positive correlation (Figure 5).

**Figure 5:**
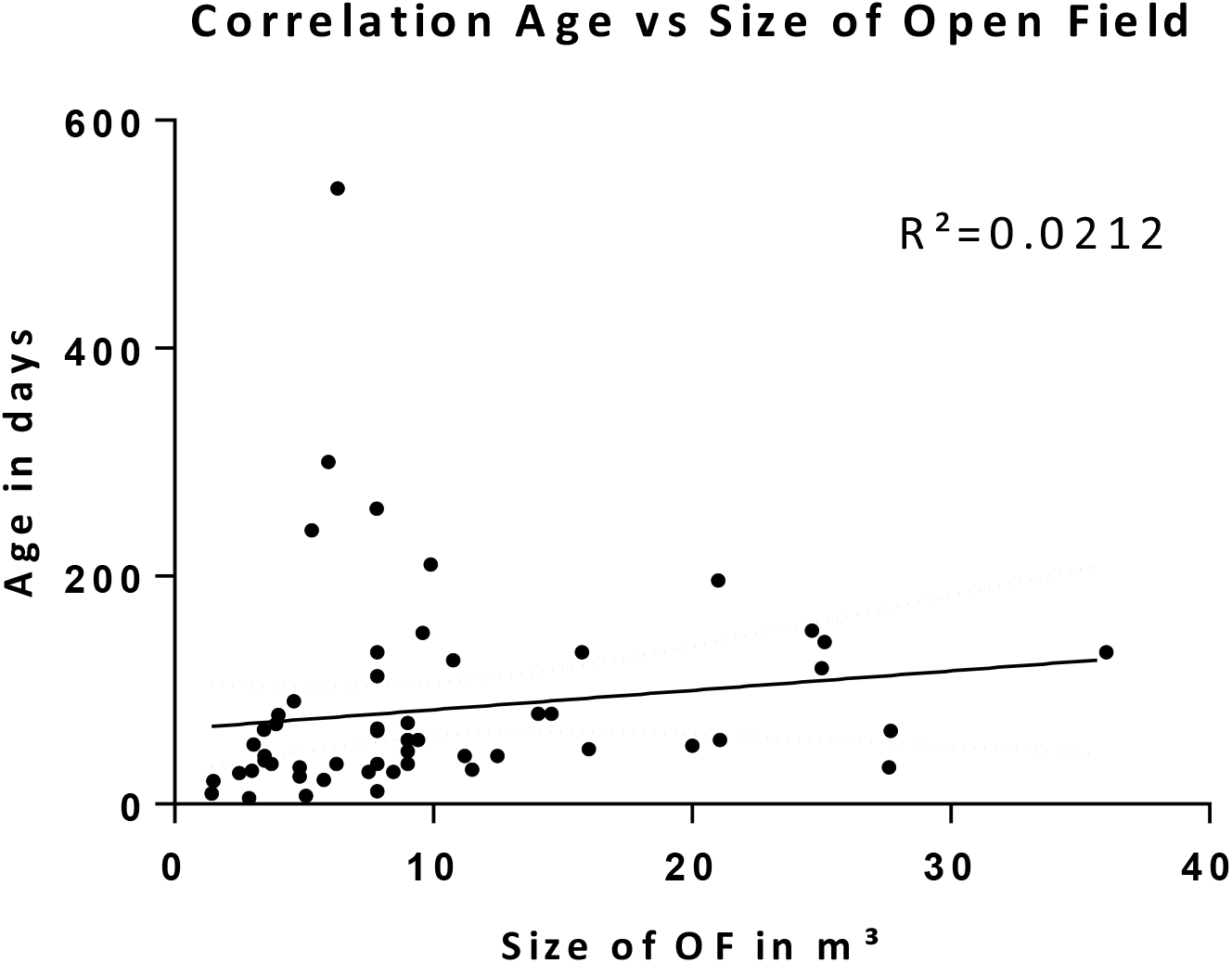
No correlation (Pearson R) between age and size of Open Field could be found (n=58).

The majority of the 24 studies provided information about the day time the test took place performed the OF before noon and 54.5% between 10 and 11am (Figure 6). Four of the studies did mention a starting time of OF at 10:00, 14:30, 15:30 and 16:30 [24, 35, 51, 75]. One study postponed the behavioral test to the evening (6 to 7pm) without further explanation [69]. Forty-five publications (65.2 %) did not refer to a daytime at all.

**Figure 6.**
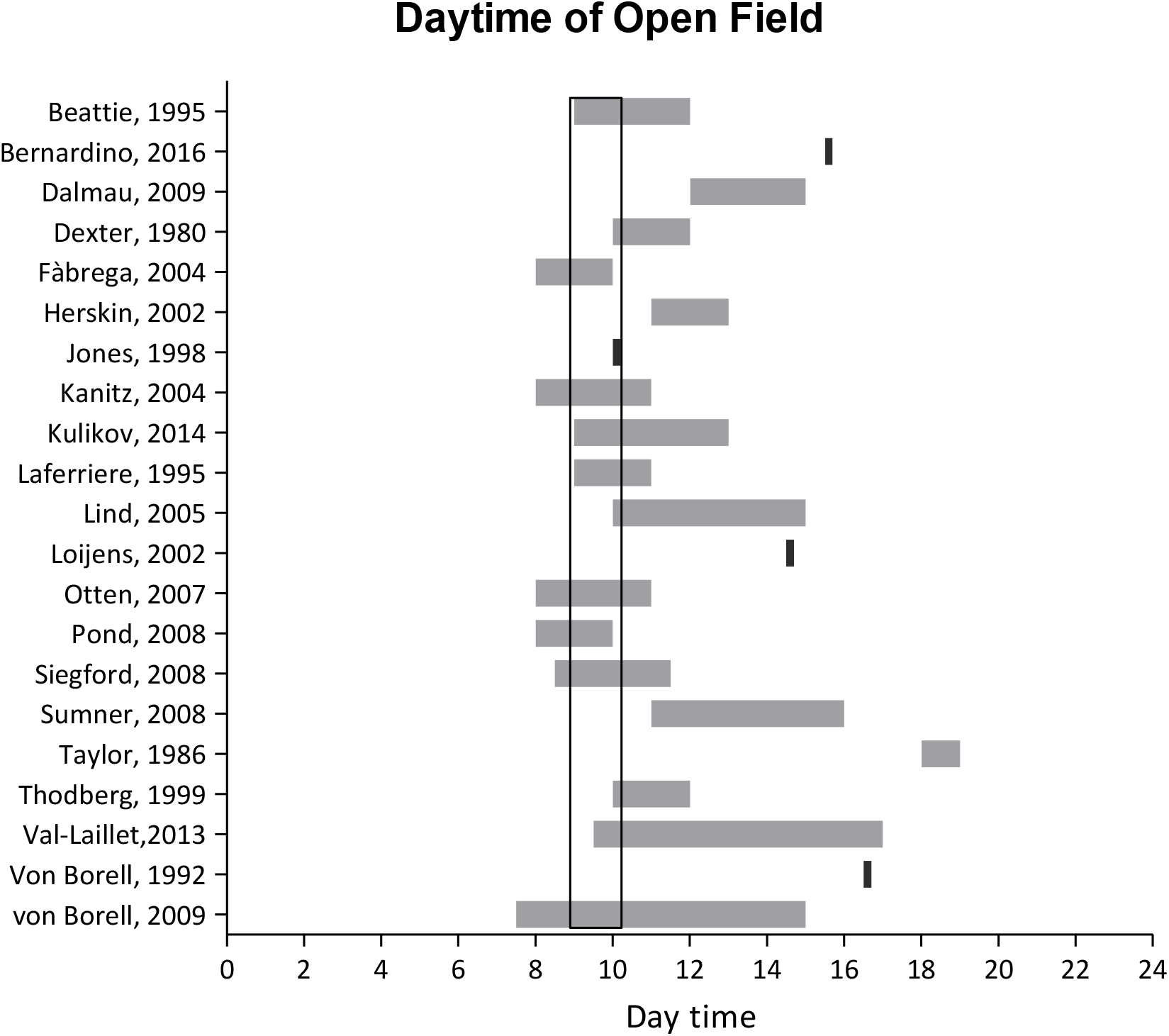
Spans of daytime the OF was performed in. Grey lines referring to the total time and black lines referring to a start time with open end.

**Figure 6:**
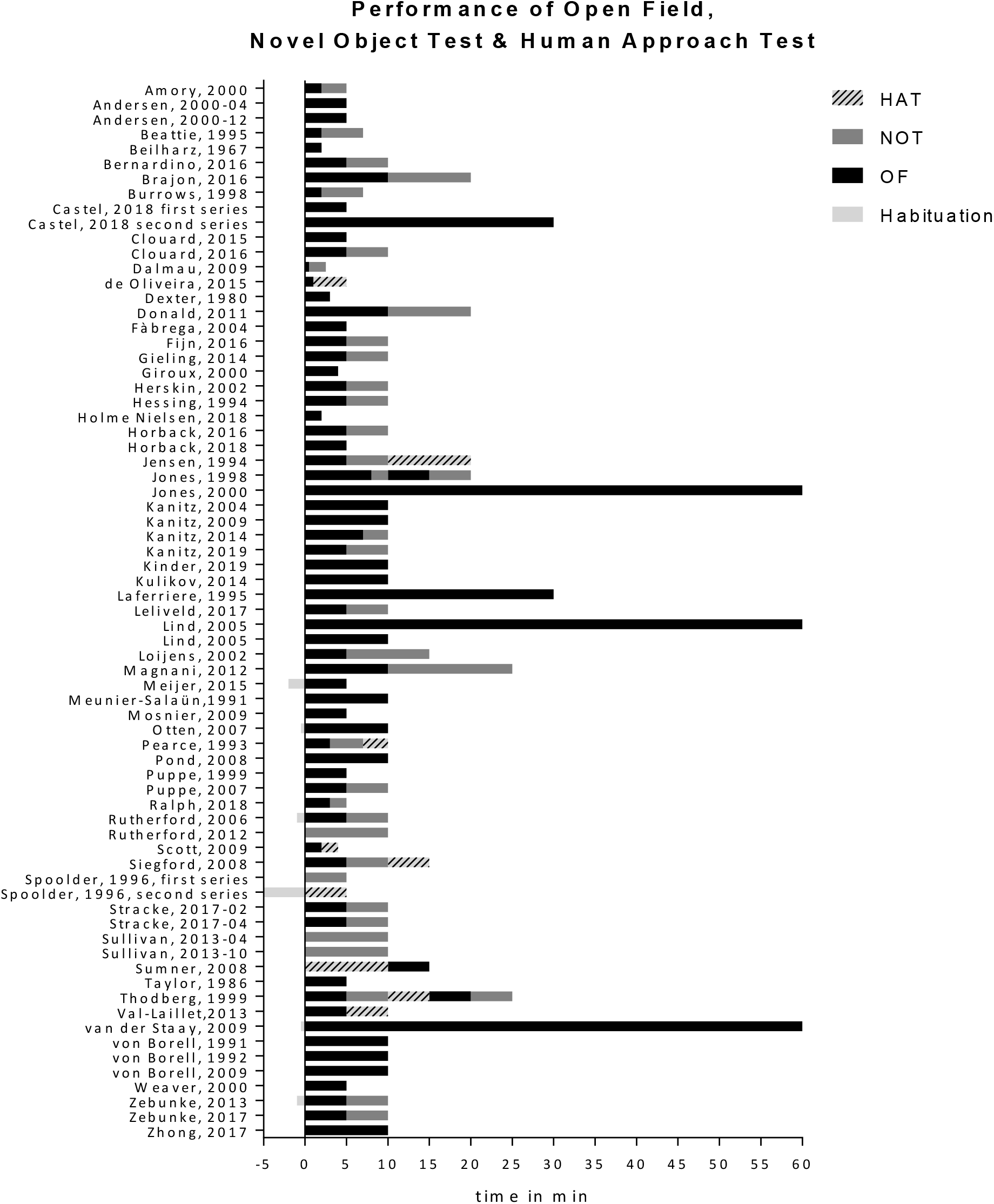
Duration pigs spent in a start box, in the Open Field with additional NOT or HAT.

A closer look at the distribution of age per sex revealed that studies using litters with both sexes mainly used younger piglets with a mean of 42.05 ± 34.1 days than the overall age of animals (Figure 7).

**Figure 7:**
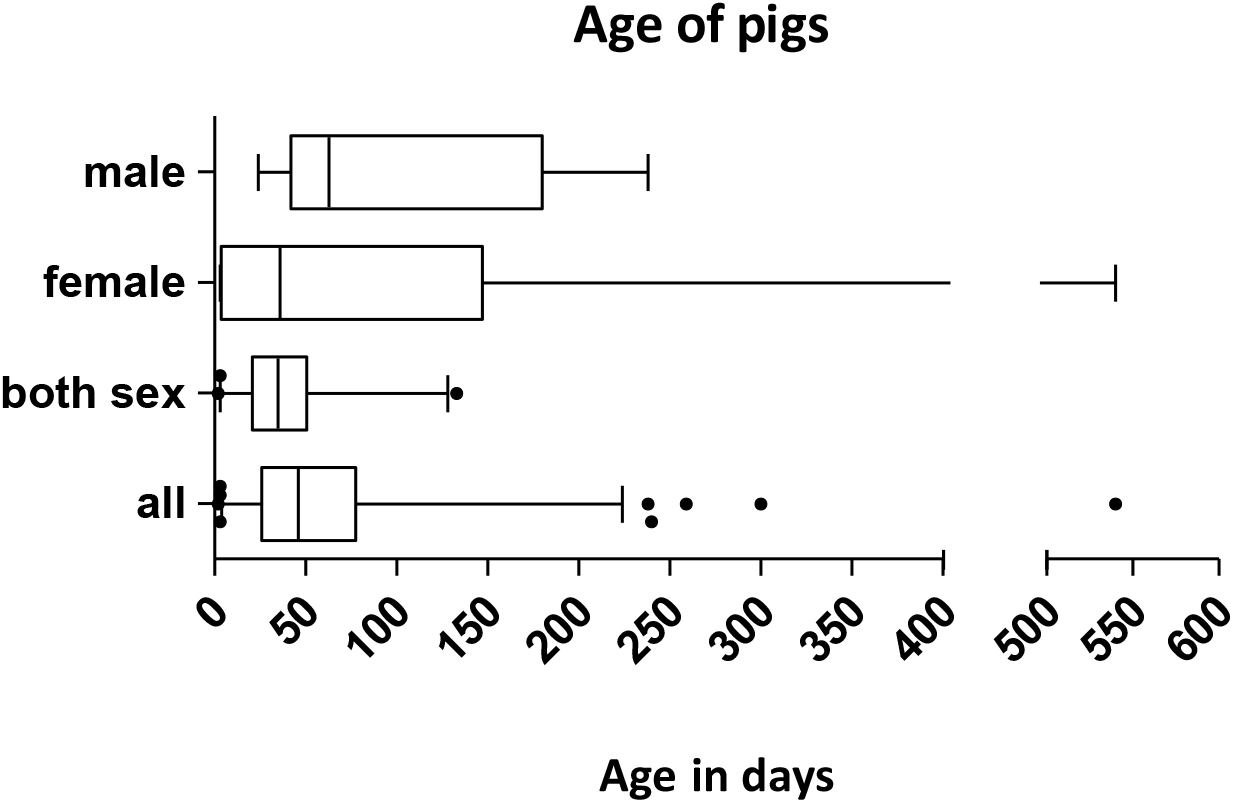
Distribution of age separated by gender.

**Figure 7:**
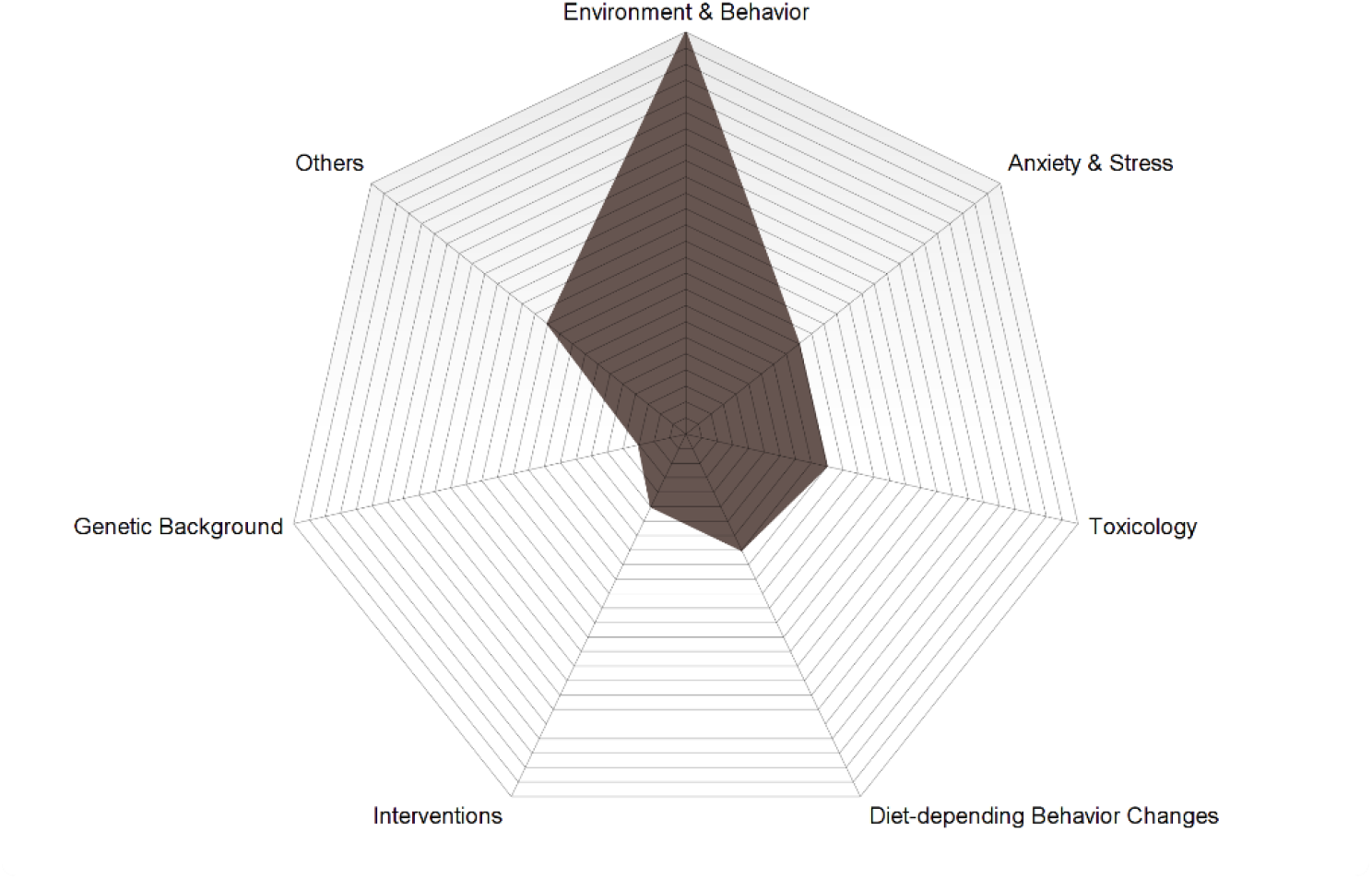
Radar plot showing the studies assigned to the categories of scientific question. Each line representing one number on the scale (n=69).

To estimate the average time a pig spent in the OF arena we distinguished between OF test, NOT and HAT. Thirty-four studies (50.7%) did only perform an OF test whereas another 35 studies additionally used NOT and/or HAT (Figure Figure 6: Duration pigs spent in a start box, in the Open Field with additional NOT or HAT.). In six studies (8.7%) a start box with an initial waiting and acclimatization time was used with a mean of 2.14min ± 1.87 [53, 62, 63, 68, 71, 77]. In the majority of studies, the OF behavioral test lasted for 10 min (8.92 min ±12.2). An intruder (human [41, 56, 62, 70, 71, 76, 92] or pig [74]) was added in eight studies. This was often combined with NOT and separated OF time. If a NOT test was used, the mean time for this test was set to 5 min (5.74 min ± 2.76).

### Parameters gained by Open Field

In this review we divided the various parameters into eight main categories and counted the number of occurrences in each publication. Those parameters only occurred less than two times where displayed as “others”. Categories in Table 1 Quality and ranking of studies based on the given information where sorted according to the frequency of occurrence.

We found mobility and immobility as well as exploratory behavior (if a NOT was used) to be the categories with the main interest followed by vocalization and excretion/ingestion. There is an overlap of parameters defined as posture in some studies with immobile parameters. Interestingly, the parameters of species-specific behavior of pigs are only ranked on the seventh position and where used in only 20 studies. If a species-specific behavior (e.g. nudging or chewing) occurred as exploratory interaction with a NOT it was categorized as exploratory behavior. Some behaviors where categorized by the authors as exploration instead of behavior. We found the wording and interpretation of parameters to be inconsistent between the various studies. Therefore, immobile parameters also describing a posture where counted in both categories.

### Domain of Scientific Question

During this systematic review as secondary outcome the question arises which experimental field are the publications assigned to, giving more insight view of how OF is used for pigs. We therefore categorized all studies given 6 main categories and cumulating the remaining ones as others.

The following categories for experimental questions were defined: Anxiety/stress, toxicology (studies of drugs and their impact on behavioral change), behavior change due to diet, genetic background, intervention (e.g. surgery), environmental change associated behavior and others (Figure 9).

### Risk of Bias

The publication bias has been applied according to the SYRCLE risk of bias tool [27] on all sixty-nine studies. Three independent authors judged all studies leading to 250 issues being solved by discussion. A full agreement on the RoB was found.

The performance bias regarding random housing concludes that no study referred to a random allocation of animals to their home pens. Mostly, animals were grouped litter wise for toleration of companions in respect to the animal welfare. However, if animals were housed in pens with the same controlled climatic conditions and sizes the author decided to answer the question positive (low risk). In this case there seems to be no negative influence of the allocation to the different pens on the experimental outcome.

Because in various publications animal behavioral tests often occurs with a special housing situation like warm areas, special signature odors or sex depended assignment, care takers or investigators could not be blinded for the experiment. In this case the authors judged against the study. If animals were assigned to all interventions with or without random allocation to the individual tests, the question of allocation concealment was defined as positive.

The complete analysis of the authors judgement with respect to the RoB is shown in Figure 10.

**Figure 10:**
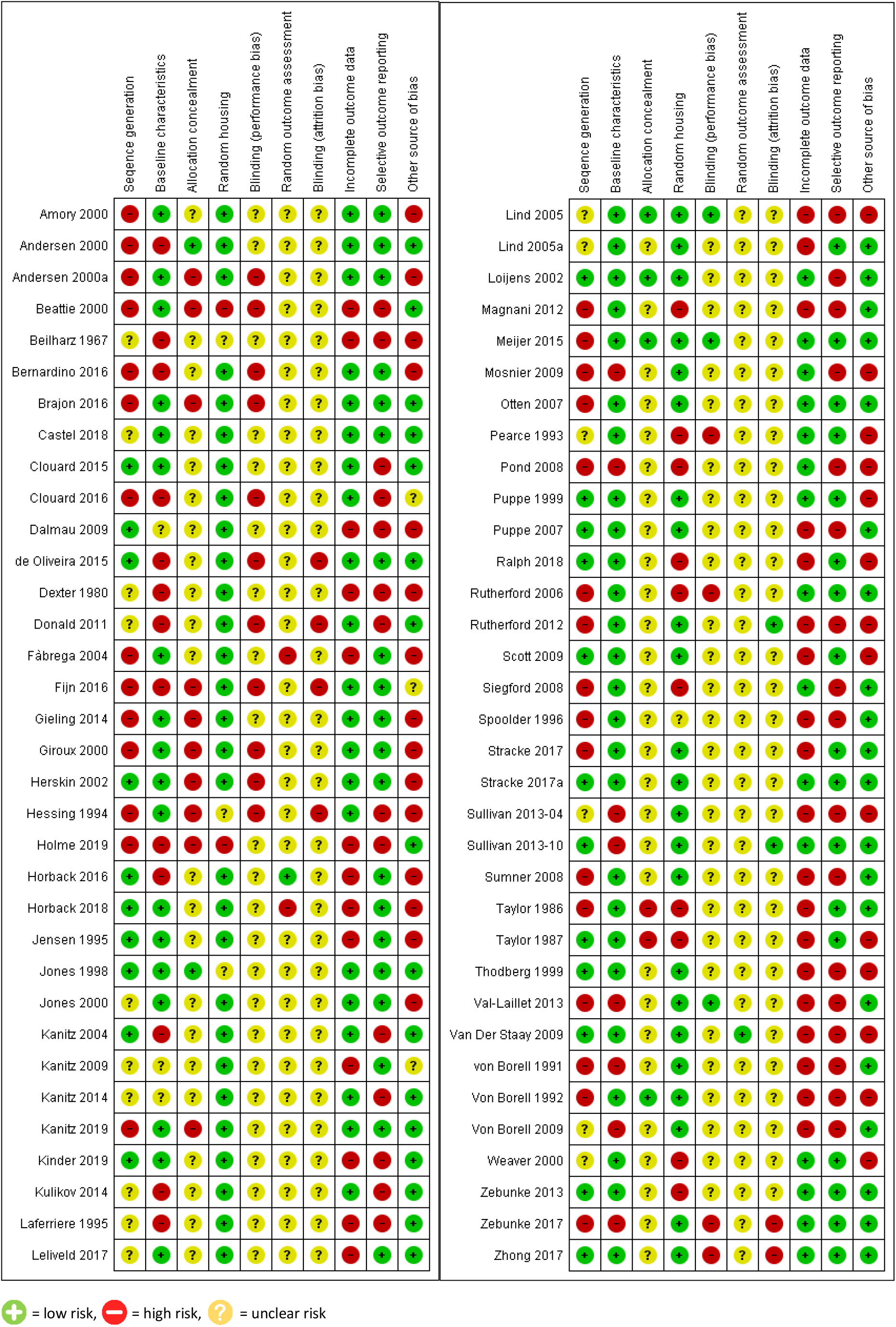
Estimated risk of bias of all studies according to the RoB-tool

Positively, around 60% of all studies had a low risk of bias regarding the baseline characteristics.

Because behavioral testing as well as grouped husbandry of pigs is highly influenced by body weight, littermates or sexes most sequence generations were not random but took this important aspect into account [84]. This leading to 30 studies with a high risk of bias in sequence generation, according to the RoB-tool.

Twenty-nine publications had a higher risk of bias mainly due to commercial funding whereas 39 studies where approved by authorities or governmental institutions leading to a low risk of bias. Additionally, we focused on the study design to answer the question of other sources of bias.

Especially, the blinding of assessors as well as the randomization of outcomes where the items with the most unclear status. Only 5 studies referred to a blinded assessor for at least one parameter.

An overview of the estimated risk of all studies per domain is shown in Figure11.

**Figure 11:**
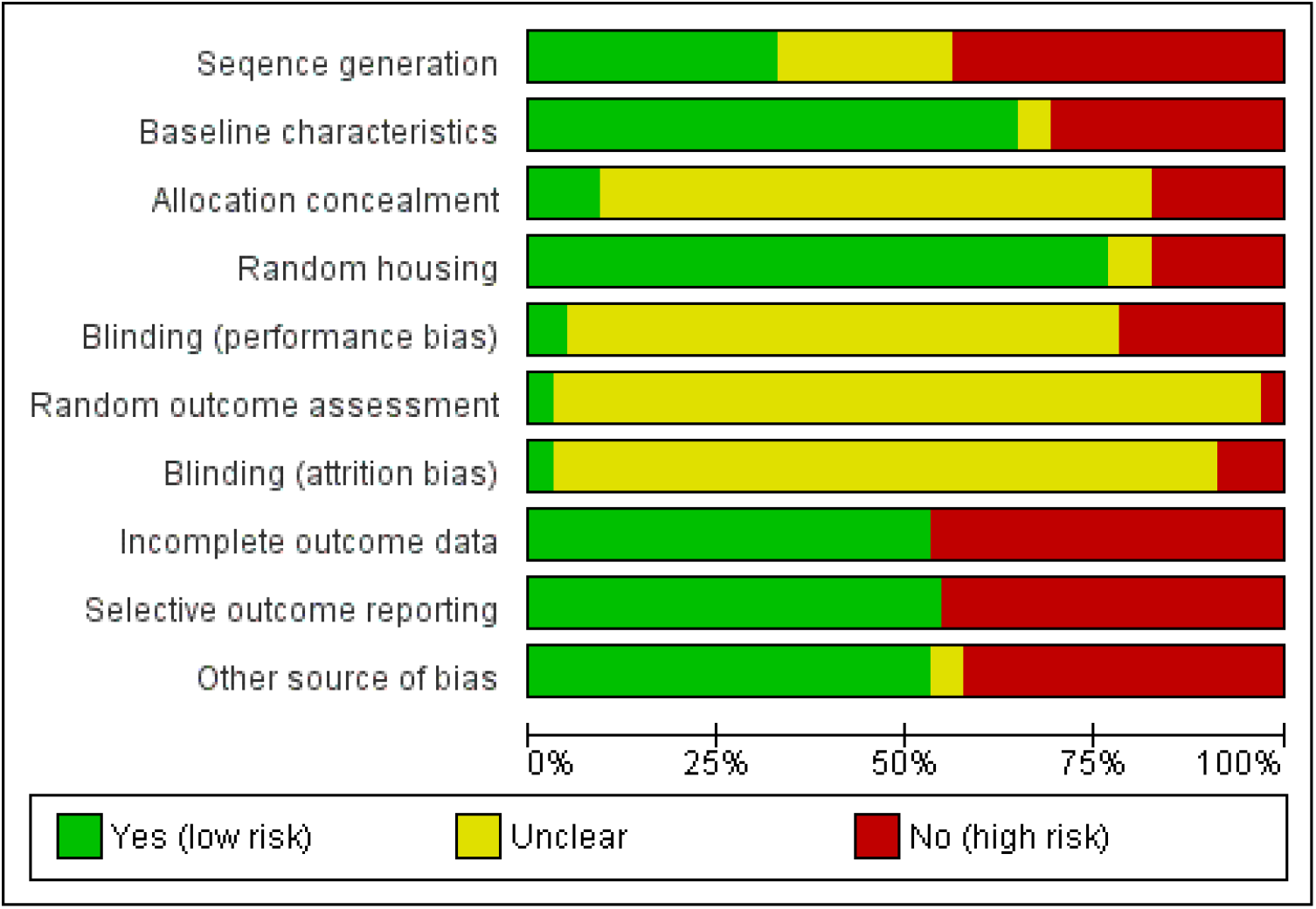
Review authors’ judgments about each risk of bias item presented as percentages across all included studies.

## Discussion

The aim of this systematic review was to investigate into the set-up of OF used in pigs. We therefore firstly looked into the size of OF arenas (Figure 2) to determine whether there is a common or standardized size being used. Although, almost half of the studies used a square or nearly-square OF the size varied widely between all studies. In mean the dimension of the OF is 11.3 m^2^. Andersen et. al referred to the size of the OF being adapted to the age of the animals [12]. In a second attempted we therefore correlated the animals age (Figure 4: Age distribution of animals per study. Age is given in days with a split x-axis at 100 days.) with the size of OF area (Figure 5). We could however not find any indications that there is a positive correlation (R^2^ = 0.02) between those parameters/outcomes. We assume the high variance might be due to structural conditions instead of being chosen in accordance of animals age or weight. In two-third of the studies the OF was subdivided into sections, whether virtually applied or painted to the floor. The comparison between the included studies showed a high variance in the number of sections from zero up to 72 (Figure 3). In median the amount of sections was 12. Overall, we found the size and number of sections of the OF to be inconsistent and not be related to the age of pigs. For future experiments, a standardization would help to compare results between studies and set them into an overall context.

Often, a male bias in research is reported where female mammals are neglected. Although in clinical research the inclusion of women reached more attention this is not the case in basic science and translational research [94–96]. In contrast to this, the distribution of gender showed multiple studies using female pigs or both sex when focusing on litters. There seems to be a low sex bias which enhances the possible translation of results to humans.

We also investigated if there is a daytime commonly used for OF tests in pigs. Rodents are nocturnal animals and most of the OF tests were performed during their inactive phase [97]. Most domestic animals are diurnal [25]. These ecological characteristics might make it difficult to translate the OF from rodents to large animals. A comparison between studies revealed the morning between 9 and 10pm to be the most used time to perform the OF test (Figure 6 Spans of daytime the OF was performed in. Grey lines referring to the total time and black lines referring to a start time with open end.). Some studies mentioned the OF were carried out after the feeding and a few used feeding as a part of the OF test [43, 88]. The predictability of food rewards or feeding are known for being stressors to rats [98]. If for anxiety test the feeding and daytime of OF test should be considered to have an effect on the outcome measures. Some studies investigated in several different tests per animal and day (data not shown). None of those seven studies performing the OF test later than 11am gave an explanation for the chosen time . Right now, no investigation into the impact of daytime on the OF could be found.

Another primary outcome of this systematic review were the test specifiers. Here we wanted to identify how long animals stayed in the OF and if there are any consistencies. Because half of the studies used the OF in combination with NOT or HAT we also wanted to picture the average time spent on these test as well. Almost half of the studies used 5 minutes and only three studies extended the time to one hour (Figure 6: Duration pigs spent in a start box, in the Open Field with additional NOT or HAT.). Therefore, the original time of 10 minutes developed by Hall [1] for rodents was often reduced down to 5 minutes. Thirty-four studies added a NOT to the OF test extending the time to 10 minutes. Of those studies, the majority investigated into the behavioral change due to environmental conditions, looking into the welfare and development of animals (Figure 7) [24, 56].

As another potential source of variability, we listed the parameters gained by the OF (Table 2). In contrast to exploratory behavior parameters of mobility especially distance and velocity are objective values which can easily be determined by using tracking software [99]. Unfortunately, many studies instead used the method of counting steps being made into sections to determine the mobility of animals. Due to different numbers of sections per area and various size, a comparison between studies is difficult, here. Additionally, the size of pigs according to their strain and age is another variable when determining the distance covered. In contrast to objective measurable values, parameters like behavior and exploration have potential limitations. The interpretation of this species-specific behavior is lacking of guidelines. Where some authors interpret the avoidance and immobility as neophobia [29] others found this to not mirror anxiety [100, 101]. Rooting and sniffing behavior changes within age of pigs decreasing over time as curiosity declines and are also be effected by the increase of test cycles [29, 42, 74]. Additionally, the husbandry conditions also influences the behavior of the pigs within the Open Field [69]. Over all, the included studies showed no consensus on the wording and categorization of parameters.

**Table 2:** Parameters gained by Open Field sorted into domain categories and their occurrence.

| Category | Description | No of occurrence/study* |
| --- | --- | --- |
| Mobility | **Total** | **113** |
|  | Distance (number of squares entered; total distance covered) | 34 |
|  | Escape attempts (by jumping) | 21 |
|  | Locomotion (without further sub-division) | 21 |
|  | Walking | 18 |
|  | Running | 7 |
|  | Trotting | 3 |
|  | Velocity | 3 |
|  | Others | 6 |
| Exploring | **Total** | **65** |
|  | Surrounding (ground, wall) | 27 |
|  | Object (sniffing/manipulating with snout or body, avoiding, looking at object, knocking object with head) | 31 |
|  | Person (touching, sniffing, looking at human, licking, biting) | 7 |
| Immobility | **Total** | **62** |
|  | Standing | 26 |
|  | Laying/Recumbency (lateral/ventral) | 10 |
|  | Time spend in center/number of entries to center | 10 |
|  | Sitting | 7 |
|  | Freezing in alert | 4 |
|  | Others | 5 |
| Excretion/ Ingestion | **Total** | **55** |
|  | Defecation | 26 |
|  | Urination | 16 |
|  | Food intake | 9 |
|  | Water intake | 4 |
| Vocalization | **Total** | **54** |
|  | Grunts | 20 |
|  | Squeals | 13 |
|  | Screams | 9 |
|  | Not differentiated (Vocalization) | 8 |
|  | Barks (Bark-like noise) | 4 |
| Posture | **Total** | **52** |
|  | Standing | 27 |
|  | Laying (ventral, lateral) | 10 |
|  | Sitting | 7 |
|  | Freezing in alert | 4 |
|  | Others | 4 |
| Behavior | **Total** | **36** |
|  | Rooting | 6 |
|  | Chewing | 5 |
|  | Sniffing | 4 |
|  | Nudging | 3 |
|  | Scratching | 3 |
|  | Watching/scanning/looking around | 3 |
|  | Others | 12 |
| Social interaction | **Total** | **13** |
|  | Aggressiveness (chasing, biting, pushing, oral grasping, attacking) | 5 |
|  | Playing with mates | 3 |
|  | Proximity to mates | 2 |
|  | Self-isolation | 2 |
|  | Exploring mates | 1 |
| Other | Study dependent parameters other than main categories | 5 |

\*multiple number of occurrence possible

A closer look into the categorization of studies to their scientific questions revealed the environment-depending behavior changes to be of most interest (Figure 7). This is followed in equal proportions of toxicological and anxiety studies. When distinguished between farm and laboratory animals it is evident that the issue of rearing and enriched environment is primarily relevant for farm animals [23, 41, 69, 85]. Other categories like surgical interventions or toxicology apply to laboratory animals. It seems the OF test is not only able to answer scientific questions which translate to humans but also a tool to assess welfare [56, 79] of farm animals. With 18 of 25 studies within this scientific field being published after the year 2000 it becomes clear that awareness for welfare of farm animals rises.

## Limitations

The quality of the included studies varied both in terms of available set-up data and comprehensible results (Table 1 Quality and ranking of studies based on the given information). The RoB analysis shows that especially within performance and detection bias the procedures were mostly unclear (Figure). We found no evidence of any study being controlled leading to a high risk of bias. Though, the sequence generation has a high risk of bias in most of the studies this does not reflect the fact that the pig as a target animal has a high sense of hierarchy. In some studies this behavior is implemented into the study question where a sequence generation cannot be random [40, 53, 60]. In other studies, it is of high interest for the pigs welfare to be grouped with equally ranked conspecifics or siblings to reduce biting or gain access to feeding [30, 60].

Our study has some inherent limitations due to the heterogeneity of data. For comparison, we recalculated the age of animals into days, which however leads to a slight difference to the actual age. Another source of bias is the limitation to English language. This unfortunately excluded some studies. A third factor of potential bias is the authors judgement of domain categories of parameters (Table 2). We tried to match and assign the different outcome parameters to their related category. All studies had a lack of information regarding the performance and detection bias. As we rely on these data, we cannot eliminate the high-risk incidental thereto. Despite these limitations, we followed the strict and sensitive protocol for data identification to reduce potential bias.

## Conclusion

There is a high need of standardization for the use of Open Field tests in pigs. Especially general terms and conditions on dimension, daytime and performance should be made. Additionally, a critical review of parameters gained by Open Field as well as terminology should be in focus. It can be stated that the use of OF alone does not give valuable information on anxiety models and is highly influenced by several external factors. In addition, apparently the number of tests and age of the animals influence objective parameters like mobility and elimination. It should be critically examined for each experimental setup whether the Open Field is a meaningful test for pigs and whether a transfer from rodent models is applicable.

## Authors contribution

MS and RT designed this study. MS and LZ searched and collected the data. MS wrote the manuscript. MK designed the graphics. MS, MK and LZ judged the RoB. All authors reviewed and revised the paper.

## References

1. Hall CS. Emotional behavior in the rat. I. Defecation and urination as measures of individual differences in emotionality. Journal of Comparative psychology. 1934;18(3):385.

2. Castanheira L, Ferreira MF, Sebastiao AM, Telles-Correia D. Anxiety Assessment in Pre-clinical Tests and in Clinical Trials: A Critical Review. Curr Top Med Chem. 2018;18(19):1656–76.

3. Tatem KS, Quinn JL, Phadke A, Yu Q, Gordish-Dressman H, Nagaraju K. Behavioral and locomotor measurements using an open field activity monitoring system for skeletal muscle diseases. J Vis Exp. 2014(91):51785.

4. Lau AA, Crawley AC, Hopwood JJ, Hemsley KM. Open field locomotor activity and anxiety-related behaviors in mucopolysaccharidosis type IIIA mice. Behav Brain Res. 2008;191(1):130–6.

5. Sturman O, Germain PL, Bohacek J. Exploratory rearing: a context- and stress-sensitive behavior recorded in the open-field test. Stress. 2018;21(5):443–52.

6. Delprato A, Algeo MP, Bonheur B, Bubier JA, Lu L, Williams RW, et al. QTL and systems genetics analysis of mouse grooming and behavioral responses to novelty in an open field. Genes Brain Behav. 2017;16(8):790–9.

7. Markowska AL, Spangler EL, Ingram DK. Behavioral assessment of the senescence-accelerated mouse (SAM P8 and R1). Physiol Behav. 1998;64(1):15–26.

8. Murphy E, Nordquist RE, van der Staay FJ. A review of behavioural methods to study emotion and mood in pigs, Sus scrofa. Applied Animal Behaviour Science. 2014;159:9–28.

9. Swindle MM, Makin A, Herron AJ, Clubb FJ, Jr., Frazier KS. Swine as models in biomedical research and toxicology testing. Vet Pathol. 2012;49(2):344–56.

10. Clouard C, Meunier-Salaun MC, Val-Laillet D. Food preferences and aversions in human health and nutrition: how can pigs help the biomedical research? Animal. 2012;6(1):118–36.

11. Sullivan S, Friess SH, Ralston J, Smith C, Propert KJ, Rapp PE, et al. Behavioral deficits and axonal injury persistence after rotational head injury are direction dependent. J Neurotrauma. 2013;30(7):538–45.

12. Andersen IL, Boe KE, Foerevik G, Janczak AM, Bakken M. Behavioural evaluation of methods for assessing fear responses in weaned pigs. Applied Animal Behaviour Science. 2000;69(3):227–40.

13. Mogil JS. Animal models of pain: progress and challenges. Nat Rev Neurosci. 2009;10(4):283–94.

14. Joshi SK, Honore P. Animal models of pain for drug discovery. Expert Opin Drug Discov. 2006;1(4):323–34.

15. Castel D, Willentz E, Doron O, Brenner O, Meilin S. Characterization of a porcine model of post-operative pain. Eur J Pain. 2014;18(4):496–505.

16. Welberg LA, Seckl JR, Holmes MC. Prenatal glucocorticoid programming of brain corticosteroid receptors and corticotrophin-releasing hormone: possible implications for behaviour. Neuroscience. 2001;104(1):71–9.

17. Lind NM, Olsen AK, Moustgaard A, Jensen SB, Jakobsen S, Hansen AK, et al. Mapping the amphetamine-evoked dopamine release in the brain of the Gottingen minipig. Brain Res Bull. 2005;65(1):1–9.

18. Geyer MA, Ellenbroek B. Animal behavior models of the mechanisms underlying antipsychotic atypicality. Prog Neuropsychopharmacol Biol Psychiatry. 2003;27(7):1071–9.

19. Winterdahl M, Noer O, Orlowski D, Schacht AC, Jakobsen S, Alstrup AKO, et al. Sucrose intake lowers mu-opioid and dopamine D2/3 receptor availability in porcine brain. Sci Rep. 2019;9(1):16918.

20. Nozari M, Shabani M, Hadadi M, Atapour N. Enriched environment prevents cognitive and motor deficits associated with postnatal MK-801 treatment. Psychopharmacology (Berl). 2014;231(22):4361–70.

21. Urakawa S, Mitsushima D, Shimozuru M, Sakuma Y, Kondo Y. An enriched rearing environment calms adult male rat sexual activity: implication for distinct serotonergic and hormonal responses to females. PLoS One. 2014;9(2):e87911.

22. Beattie VE, Walker N, Sneddon IA. Effect of rearing environment and change of environment on the behaviour of gilts. Applied Animal Behaviour Science. 1995;46(1-2):57–65.

23. Fijn L, Antonides A, Aalderink D, Nordquist RE, van der Staay FJ. Does litter size affect emotionality, spatial learning and memory in piglets? Applied Animal Behaviour Science. 2016;178:23–31.

24. Jones R, Nicol CJ. A note on the effect of control of the thermal environment on the well-being of growing pigs. Applied Animal Behaviour Science. 1998;60(1):1–9.

25. Forkman B, Boissy A, Meunier-Salaün MC, Canali E, Jones RB. A critical review of fear tests used on cattle, pigs, sheep, poultry and horses. Physiology and Behavior. 2007;92(3):340–74.

26. Moher D, Liberati A, Tetzlaff J, Altman DG. Preferred reporting items for systematic reviews and meta-analyses: the PRISMA statement. PLoS Med. 2009;6(7):e1000097.

27. Hooijmans CR, Rovers MM, de Vries RB, Leenaars M, Ritskes-Hoitinga M, Langendam MW. SYRCLE’s risk of bias tool for animal studies. BMC Med Res Methodol. 2014;14:43.

28. Higgins JP, Altman DG, Gotzsche PC, Juni P, Moher D, Oxman AD, et al. The Cochrane Collaboration’s tool for assessing risk of bias in randomised trials. Bmj. 2011;343:d5928.

29. Donald RD, Healy SD, Lawrence AB, Rutherford KMD. Emotionality in growing pigs: Is the open field a valid test? Physiology & Behavior. 2011;104(5):906–13.

30. Giroux S, Martineau GP, Robert S. Relationships between individual behavioural traits and post-weaning growth in segregated early-weaned piglets. Applied Animal Behaviour Science. 2000;70(1):41–8.

31. Lind NM, Vinther M, Hemmingsen RP, Hansen AK. Validation of a digital video tracking system for recording pig locomotor behaviour. J Neurosci Methods. 2005;143(2):123–32.

32. Hessing MJC, Hagelso AM, Schouten WGP, Wiepkema PR, Vanbeek JAM. INDIVIDUAL BEHAVIORAL AND PHYSIOLOGICAL STRATEGIES IN PIGS. Physiology & Behavior. 1994;55(1):39–46.

33. Von Borell E, Ladewig J. RELATIONSHIP BETWEEN BEHAVIOR AND ADRENOCORTICAL-RESPONSE PATTERN IN DOMESTIC PIGS. Applied Animal Behaviour Science. 1992;34(3):195–206.

34. Horback KM, Parsons TD. Temporal stability of personality traits in group-housed gestating sows. Animal. 2016;10(8):1351–9.

35. Bernardino T, Tatemoto P, Morrone B, Rodrigues PHM, Zanella AJ. Piglets born from sows fed high fibre diets during pregnancy are less aggressive prior to weaning. PLoS ONE. 2016;11(12).

36. Jensen KH, Oksbjerg N, Jørgensen E. Dietary salbutamol and level of protein: effects on the acute stress response in pigs. Physiol Behav. 1994;55(2):375–9.

37. Castel D, Sabbag I, Nasaev E, Peng S, Meilin S. Open field and a behavior score in PNT model for neuropathic pain in pigs. J Pain Res. 2018;11:2279–93.

38. Jones JB, Wathes CM, White RP, Jones RB. Do pigs find a familiar odourant attractive in novel surroundings? Applied Animal Behaviour Science. 2000;70(2):115–26.

39. Clouard C, Gerrits WJJ, Kemp B, Val-Laillet D, Bolhuis JE. Perinatal Exposure to a Diet High in Saturated Fat, Refined Sugar and Cholesterol Affects Behaviour, Growth, and Feed Intake in Weaned Piglets. Plos One. 2016;11(5).

40. Kanitz E, Hameister T, Tuchscherer M, Tuchscherer A, Puppe B. Social support attenuates the adverse consequences of social deprivation stress in domestic piglets. Horm Behav. 2014;65(3):203–10.

41. de Oliveira D, da Costa M, Zupan M, Rehn T, Keeling LJ. Early human handling in non-weaned piglets: Effects on behaviour and body weight. Applied Animal Behaviour Science. 2015;164:56–63.

42. Kanitz E, Puppe B, Tuchscherer M, Heberer M, Viergutz T, Tuchscherer A. A single exposure to social isolation in domestic piglets activates behavioural arousal, neuroendocrine stress hormones, and stress-related gene expression in the brain. Physiology & Behavior. 2009;98(1-2):176–85.

43. Fàbrega E, Diestre A, Font J, Carrión D, Velarde A, Ruiz-de-la-Torre JL, et al. Differences in open field behavior between heterozygous and homozygous negative gilts for the RYR(1) gene. Journal of Applied Animal Welfare Science. 2004;7(2):83–93.

44. Kanitz E, Tuchscherer M, Otten W, Tuchscherer A, Zebunke M, Puppe B. Coping Style of Pigs Is Associated With Different Behavioral, Neurobiological and Immune Responses to Stressful Challenges. Front Behav Neurosci. 2019;13:173.

45. Kinder HA, Baker EW, Howerth EW, Duberstein KJ, West FD. Controlled Cortical Impact Leads to Cognitive and Motor Function Deficits that Correspond to Cellular Pathology in a Piglet Traumatic Brain Injury Model. J Neurotrauma. 2019;36(19):2810–26.

46. Gieling ET, Antonides A, Fink-Gremmels J, Ter Haar K, Kuller WI, Meijer E, et al. Chronic allopurinol treatment during the last trimester of pregnancy in sows: effects on low and normal birth weight offspring. PLoS One. 2014;9(1):e86396.

47. Kulikov VA, Khotskin NV, Nikitin SV, Lankin VS, Kulikov AV, Trapezov OV. Application of 3-D imaging sensor for tracking minipigs in the open field test. Journal of Neuroscience Methods. 2014;235:219–25.

48. Herskin MS, Jensen KH. Effects of open field testing and associated handling v. handling alone on the adrenocortical reactivity of piglets around weaning. Animal Science. 2002;74:485–91.

49. Lind NM, Arnfred SM, Hemmingsen RP, Hansen K, Jensen KH. Open field behaviour and reaction to novelty in Gottingen minipigs: Effects of amphetamine and haloperidol. Scandinavian Journal of Laboratory Animal Science. 2005;32(2):103–12.

50. Holme Nielsen C, Bladt Brandt A, Thymann T, Obelitz-Ryom K, Jiang P, Vanden Hole C, et al. Rapid Postnatal Adaptation of Neurodevelopment in Pigs Born Late Preterm. Developmental Neuroscience. 2019;40(5-6):586–600.

51. Loijens LWS, Schouten WGP, Wiepkema PR, Wiegant VM. Brain opioid receptor density reflects behavioral and heart rate responses in pigs. Physiology & Behavior. 2002;76(4-5):579–87.

52. Horback KM, Parsons TD. Ontogeny of behavioral traits in commercial sows. Animal. 2018;12(11):2365–72.

53. Magnani D, Cafazzo S, Calà P, Costa LN. Searching for differences in the behavioural response of piglet groups subjected to novel situations. Behavioural Processes. 2012;89(1):68–73.

54. Meuniersalaun MC, Monnier M, Colleaux Y, Seve B, Henry Y. IMPACT OF DIETARY TRYPTOPHAN AND BEHAVIORAL TYPE ON BEHAVIOR, PLASMA-CORTISOL, AND BRAIN-METABOLITES OF YOUNG-PIGS. Journal of Animal Science. 1991;69(9):3689–98.

55. Kanitz E, Tuchscherer M, Puppe B, Tuchscherer A, Stabenow B. Consequences of repeated early isolation in domestic piglets (Sus scrofa) on their behavioural, neuroendocrine, and immunological responses. Brain Behav Immun. 2004;18(1):35–45.

56. Pearce GP, Paterson AM. THE EFFECT OF SPACE RESTRICTION AND PROVISION OF TOYS DURING REARING ON THE BEHAVIOR, PRODUCTIVITY AND PHYSIOLOGY OF MALE PIGS. Applied Animal Behaviour Science. 1993;36(1):11–28.

57. Laferrière A, Ertug F, Moss IR. Prenatal cocaine alters open-field behavior in young swine. Neurotoxicol Teratol. 1995;17(2):81–7.

58. Rutherford KMD, Donald RD, Lawrence AB, Wemelsfelder F. Qualitative Behavioural Assessment of emotionality in pigs. Applied Animal Behaviour Science. 2012;139(3-4):218–24.

59. Leliveld LMG, Dupjan S, Tuchscherer A, Puppe B. Vocal correlates of emotional reactivity within and across contexts in domestic pigs (Sus scrofa). Physiology & Behavior. 2017;181:117–26.

60. Rutherford KMD, Haskell MJ, Glasbey C, Lawrence AB. The responses of growing pigs to a chronic-intermittent stress treatment. Physiology & Behavior. 2006;89(5):670–80.

61. Meijer E, van Nes A, Back W, van der Staay FJ. Clinical effects of buprenorphine on open field behaviour and gait symmetry in healthy and lame weaned piglets. Vet J. 2015;206(3):298–303.

62. Spoolder HAM, Burbidge JA, Lawrence AB, Simmins PH, Edwards SA. Individual behavioural differences in pigs: Intra- and inter-test consistency. Applied Animal Behaviour Science. 1996;49(2):185–98.

63. Mosnier E, Dourmad JY, Etienne M, Le Floc’h N, Pere MC, Ramaekers P, et al. Feed intake in the multiparous lactating sow: Its relationship with reactivity during gestation and tryptophan status. Journal of Animal Science. 2009;87(4):1282–91.

64. Stracke J, Otten W, Tuchscherer A, Puppe B, Dupjan S. Serotonin depletion induces pessimistic-like behavior in a cognitive bias paradigm in pigs. Physiology & Behavior. 2017;174:18–26.

65. Otten W, Kanitz E, Tuchscherer M, Puppe B, Nurnberg G. Repeated administrations of adrenocorticotropic hormone during gestation in gilts: Effects on growth, behaviour and immune responses of their piglets. Livestock Science. 2007;106(2-3):261–70.

66. Stracke J, Otten W, Tuchscherer A, Witthahn M, Metges CC, Puppe B, et al. Dietary tryptophan supplementation and affective state in pigs. Journal of Veterinary Behavior-Clinical Applications and Research. 2017;20:82–90.

67. Pond WG, Mersmann HJ, Su DR, McGlone JJ, Wheeler MB, Smith EO. Neonatal dietary cholesterol and Alleles of cholesterol 7-alpha hydroxylase affect piglet cerebrum weight, cholesterol concentration and behavior. Journal of Nutrition. 2008;138(2):282–6.

68. Ralph C, Hebart M, Cronin GM. Enrichment in the Sucker and Weaner Phase Altered the Performance of Pigs in Three Behavioural Tests. Animals (Basel). 2018;8(5).

69. Taylor L, Friend TH. OPEN-FIELD TEST BEHAVIOR OF GROWING SWINE MAINTAINED ON A CONCRETE FLOOR AND A PASTURE. Applied Animal Behaviour Science. 1986;16(2):143–8.

70. Siegford JM, Rucker G, Zanella AJ. Effects of pre-weaning exposure to a maze on stress responses in pigs at weaning and on subsequent performance in spatial and fear-related tests. Applied Animal Behaviour Science. 2008;110(1-2):189–202.

71. Val-Laillet D, Tallet C, Guerin C, Meunier-Salaun MC. Behavioural reactivity, social and cognitive abilities of Vietnamese and Pitman-Moore weaned piglets. Applied Animal Behaviour Science. 2013;148(1-2):108–19.

72. Sullivan S, Friess SH, Ralston J, Smith C, Propert KJ, Rapp PE, et al. Improved behavior, motor, and cognition assessments in neonatal piglets. J Neurotrauma. 2013;30(20):1770–9.

73. Van Der Staay FJ, Pouzet B, Mahieu M, Nordquist RE, Schuurman T. The d-amphetamine-treated Göttingen miniature pig: An animal model for assessing behavioral effects of antipsychotics. Psychopharmacology. 2009;206(4):715–29.

74. Sumner BEH, D’Eath RB, Farnworth MJ, Robson S, Russell JA, Lawrence AB, et al. Early weaning results in less active behaviour, accompanied by lower 5-HT1A and higher 5-HT2A receptor mRNA expression in specific brain regions of female pigs. Psychoneuroendocrinology. 2008;33(8):1077–92.

75. von Borell E, Hurnik JF. Stereotypic behavior, adrenocortical function, and open field behavior of individually confined gestating sows. Physiol Behav. 1991;49(4):709–13.

76. Thodberg K, Jensen KH, Herskin MS. A general reaction pattern across situations in prepubertal gilts. Applied Animal Behaviour Science. 1999;63(2):103–19.

77. Weaver SA, Aherne FX, Meaney MJ, Schaefer AL, Dixon WT. Neonatal handling permanently alters hypothalamic-pituitary-adrenal axis function, behaviour, and body weight in boars. J Endocrinol. 2000;164(3):349–59.

78. Zhong M, Shoemake C, Fuller A, White D, Hanks C, Brocksmith D, et al. Development of a Functional Observational Battery in the Minipig for Regulatory Neurotoxicity Assessments. International Journal of Toxicology. 2017;36(2):113–23.

79. Zebunke M, Puppe B, Langbein J. Effects of cognitive enrichment on behavioural and physiological reactions of pigs. Physiology & Behavior. 2013;118:70–9.

80. Amory JR, Pearce GP. Alarm pheromones in urine modify the behaviour of weaner pigs. Animal Welfare. 2000;9(2):167–75.

81. Andersen IL, Faerevik G, Boe KE, Janczak AM, Bakken M. Effects of diazepam on the behaviour of weaned pigs in three putative models of anxiety. Applied Animal Behaviour Science. 2000;68(2):121–30.

82. Beilharz RG, Cox DF. Genetic analysis of open field behavior in swine. J Anim Sci. 1967;26(5):988–90.

83. Puppe B, Ernst K, Schon PC, Manteuffel G. Cognitive enrichment affects behavioural reactivity in domestic pigs. Applied Animal Behaviour Science. 2007;105(1-3):75–86.

84. Brajon S, Laforest JP, Schmitt O, Devillers N. A preliminary study of the effects of individual response to challenge tests and stress induced by humans on learning performance of weaned piglets (Sus scrofa). Behavioural Processes. 2016;129:27–36.

85. Taylor L, Friend TH. EFFECT OF HOUSING ON OPEN-FIELD TEST BEHAVIOR OF GESTATING GILTS. Applied Animal Behaviour Science. 1987;17(1-2):83–93.

86. Clouard C, Gerrits WJJ, van Kerkhof I, Smink W, Bolhuis JE. Dietary Linoleic and a-Linolenic Acids Affect Anxiety-Related Responses and Exploratory Activity in Growing Pigs. Journal of Nutrition. 2015;145(2):358–64.

87. von Borell E, Bunger B, Schmidt T, Horn T. Vocal-type classification as a tool to identify stress in piglets under on-farm conditions. Animal Welfare. 2009;18(4):407–16.

88. Dalmau A, Fabrega E, Velarde A. Fear assessment in pigs exposed to a novel object test. Applied Animal Behaviour Science. 2009;117(3-4):173–80.

89. Zebunke M, Nurnberg G, Melzer N, Puppe B. The backtest in pigs revisited-Inter-situational behaviour and animal classification. Applied Animal Behaviour Science. 2017;194:7–13.

90. Dexter JD, Tumbleson ME, Decker JD, Middleton CC. Fetal alcohol syndrome in Sinclair (S-1) miniature swine. Alcoholism: Clinical and Experimental Research. 1980;4(2):146–51.

91. Puppe B, Schon PC, Wendland K. Monitoring of piglets’ open field activity and choice behaviour during the replay of maternal vocalization: a comparison between Observer and PID technique. Laboratory Animals. 1999;33(3):215–20.

92. Scott K, Laws DM, Courboulay V, Meunier-Salaun MC, Edwards SA. Comparison of methods to assess fear of humans in sows. Applied Animal Behaviour Science. 2009;118(1-2):36–41.

93. Margulies S, Sullivan S, Friess S, Ralston J, Smith C, Propert K, et al. Behavior, motor, and cognition assessments in neonatal piglets. Journal of Neurotrauma. 2012;29(10):A214–A5.

94. Yoon DY, Mansukhani NA, Stubbs VC, Helenowski IB, Woodruff TK, Kibbe MR. Sex bias exists in basic science and translational surgical research. Surgery. 2014;156(3):508–16.

95. Beery AK, Zucker I. Sex bias in neuroscience and biomedical research. Neuroscience and biobehavioral reviews. 2011;35(3):565–72.

96. Howard LM, Ehrlich AM, Gamlen F, Oram S. Gender-neutral mental health research is sex and gender biased. The lancet Psychiatry. 2017;4(1):9–11.

97. Datta S, Samanta D, Tiwary B, Chaudhuri AG, Chakrabarti N. Sex and estrous cycle dependent changes in locomotor activity, anxiety and memory performance in aged mice after exposure of light at night. Behavioural brain research. 2019;365:198–209.

98. Tazi A, Dantzer R, Le Moal M. Prediction and control of food rewards modulate endogenous pain inhibitory systems. Behav Brain Res. 1987;23(3):197–204.

99. Zhang C, Li H, Han R. An open-source video tracking system for mouse locomotor activity analysis. BMC Res Notes. 2020;13(1):48.

100. Jensen P, Forkman B, Thodberg K, Koster E. INDIVIDUAL VARIATION AND CONSISTENCY IN PIGLET BEHAVIOR. Applied Animal Behaviour Science. 1995;45(1-2):43–52.

101. Prut L, Belzung C. The open field as a paradigm to measure the effects of drugs on anxiety-like behaviors: a review. Eur J Pharmacol. 2003;463(1-3):3–33.

